# A rapid, sensitive, and quantitative high plex biomarker digital detection platform enabled by Hypercoding

**DOI:** 10.64898/2026.03.23.711448

**Authors:** Madhavi Bathina, Angela P. Blum, Jeff Brodin, Nicole DeBuono, Yao Fu, Bo Lu, Matthew R. Naticchia, Daniel Ortiz, Andrew Richards, Cyrus de Rozières, Rachel Schowalter, Sarah Shultzaberger, Samantha Snow, Stephen Tanner, Christy L. Trejo, Spencer Ward, Jaekyung Koh, Richard LeCoultre, Kristen Read, Shashank Sathe, Christian Schlegel, Isabelle Schlegel, Sebastian Shaner, James Tsay, Jessica Weir, Ka Man Wong, Kameel Abi-Samra, Julia Alldredge, Paul Anderson, John Bailey, Celyne Bollig, Julie Bonnardel, Adrien Bru, Antony C. S. Chan, Tim Chang, Lisa DeBerg, John van Doorn, PJ Driscoll, Travis Duarte, Alexander Esparza, Drew Frerichs, Alain Gautherot, Lance Held, Garren Hendricks, Greg Holst, Kelli Iwamoto, Hector Jimenez, Omid Khandan, Stephen Kleppinger, Brenndon Lee, Panfeng Li, Donald Maine, Manuel Mendez, Kei Miyagi, Greg Nagy, Alex Nguyen, David Oaks, Deepak Rawat, Berard Schultheis, Dylan Stalcup, Gavin Stone, Michael J. Stone, Scott Sylliaasen, Eileen Thai, Jason Trachewsky, Chuong Tran, Tom Uhl, David Vigil, Jeremy Ward, Erik Williamson, Roshan Yohannan, Pieter van Rooyen, Vikram Vaz, Lorenzo Berti, Ludovic G. Vincent

**Author notes:** **Corresponding Author:** Ludovic Vincent, Pleno Inc., 6275 Nancy Ridge Dr., Suite 100, San Diego, CA, 92121. **Authorship**: See <u>Acknowledgements</u> section for author contributions.

## Abstract

Low-cost, multiplexed, and automated assays are needed to make omic technologies more broadly accessible in clinical, research, and commercial settings. We present Hypercoding™, a scalable technology for detection and quantitation of multi-omic targets. Drawing from data reliability methods in the telecommunications field, Hypercoding uses fluorescent signals from hybridization with an error-correcting code to enable detection of high-plexity targets from biological samples, such as human DNA. In the presence of the target, a linear DNA construct is circularized, immobilized, and amplified to enable single-molecule detection of a target via rapid readout cycles within a 96-well plate. We demonstrate capability for >10,000 code plexity and accurate (98.7%) genotyping of 209 pharmacogenomic variants. Furthermore, we show computation of copy number variation with whole chromosome and sub-gene resolution, as well as quantitation of target abundance down to 10 fM sensitivity with a dynamic range of up to 10 logs.

## Introduction

Advances in genomic and multi-omic technologies have transformed translational research and are now poised to do the same for clinical care. Genomic and multi-omic testing enable higher precision in clinical decision-making including assessing disease risk and disease diagnosis, informing drug selection, and monitoring treatment response and disease recurrence resulting in highly personalized patient care^1,2^.

To unlock the benefits of omic testing at a population level, large-scale testing is critical for both translational studies and for enabling routine adoption in the health care system^3^. Powering large-scale testing will require molecular assays that are fast, affordable, and easy to implement and standardize across many labs^4^. Next-generation sequencing (NGS) has been critical to the discovery of new clinically relevant variants and bringing new applications into the translational and clinical space. However, large-scale implementation of NGS in the clinic remains challenging due to the overall workflow complexity and long turnaround times, the need for specialized skill sets, and costly data storage requirements^5,6^. Quantitative and digital polymerase chain reaction (qPCR & dPCR) testing remains a workhorse in clinical labs due to the workflow simplicity, speed, and low cost^7,8^, but their utilities are limited to <20 targets in a single reaction given the difficulty in differentiating numerous fluorophores with standard optical detection systems, thus limiting the scalability of these approaches^9-11^.

Higher plexities can be achieved with encoded assays that link an analyte to a pre-determined classification signature that serves as a proxy readout^12-16^. Although this has been demonstrated with microarray and sequencing readouts, there is no integrated platform for fast, highly quantitative detection of proxy readouts. Here, we present a platform that enables rapid, high-plex detection and quantitation by linking analyte detection to a Hypercode™: a set of concatenated DNA segments that are detected sequentially via hybridization to fluorescently tagged detection oligo complexes, which reveals the classification signature associated with the target analyte. We further improve classification robustness by incorporating a machine learning algorithm and demonstrate the accuracy of this approach on human-derived cell-lines across 227 pharmacogenomic targets spanning a range of content types including single nucleotide variants (SNVs), insertions-deletions (indels), copy number variants (CNVs), and challenging genomic regions of high sequence homology and high GC content. Further, we demonstrate scalability to over 10,000 unique Hypercodes and measurements of absolute analyte concentrations spanning up to 10 logs of dynamic range with sensitivities in the femtomolar range.

## Results

Hypercoding of omics targets is enabled by a multi-functional DNA probe called a Plenoid™ (Methods). A Plenoid is a circularizable padlock probe^17^ that includes target recognition elements, referred to as probe arms, and a Hypercode constructed from several concatenated short identification sequences termed segments. In the presence of the proper complementary genomic variant or other analyte, Plenoids hybridize to their intended targets, and are subsequently circularized via ligation. Non-circularized Plenoids are enzymatically digested. Circularized Plenoids are then immobilized on a surface and amplified through rolling circle amplification (RCA)^14,18^ into rolling circle products (RCPs) containing multiple copies of the associated Hypercode for robust fluorescent detection (Fig. 1a, Methods).

**Figure 1.**
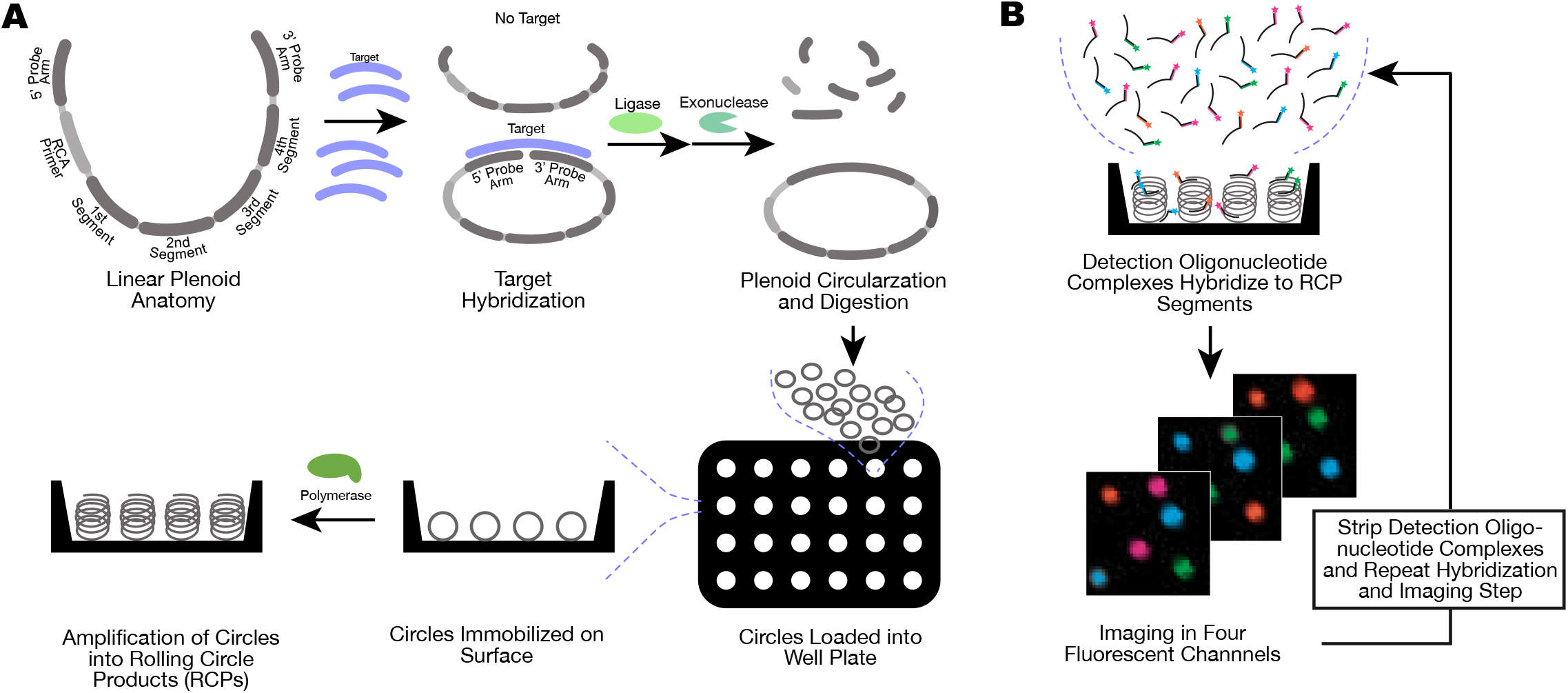
Schematic representation of Pleno assay workflow to detect omic targets. A) Formation of rolling circle products (RCPs) from a linear Plenoid. A Plenoid is a bio-engineered linear DNA construct containing a pair of target-specific probes, a set of fluorescent-probe recognition sequences termed segments, and a rolling circle amplification (RCA) primer. The first step in the Plenoid conversion assay is hybridization of target molecules to a set of Plenoids. Plenoids that do not hybridize an intended target will remain as linear DNA. In the presence of the specific target, both ends of the Plenoid are bought into proximity and a high-fidelity ligase circularizes the Plenoid. Each ligation event stems from a unique Plenoid-target binding event. Non-circularized Plenoids are subsequently enzymatically digested. Circularized Plenoids are transferred to a 96 well plate and bind to the polymer-functionalized glass surface. Plenoids are then amplified into RCPs via RCA to generate a plurality of signature Hypercode sequences in a discrete space. B) Fluorescent detection of RCPs. Segments from RCPs are probed using a set of detection oligonucleotide complexes in repeated readout cycles. After washing away unbound detection oligonucleotide complexes, each well is imaged using a four-color optical fluorescent system and images further processed to identify RCPs. To probe each Plenoid segment, multiple cycles are performed by stripping the bound detection oligonucleotide complexes and repeating the probing step with a different detection oligonucleotide complex.

To identify the unique Hypercode from individual RCPs, each segment was probed twice with a unique set of detection oligonucleotide complexes and imaged (Fig. 1b). Subsequent image analysis generated an intensity vector per object that is then matched to the nearest consensus Hypercode profile uniquely associated with its target analyte (Fig. 2a, Methods).

**Figure 2.**
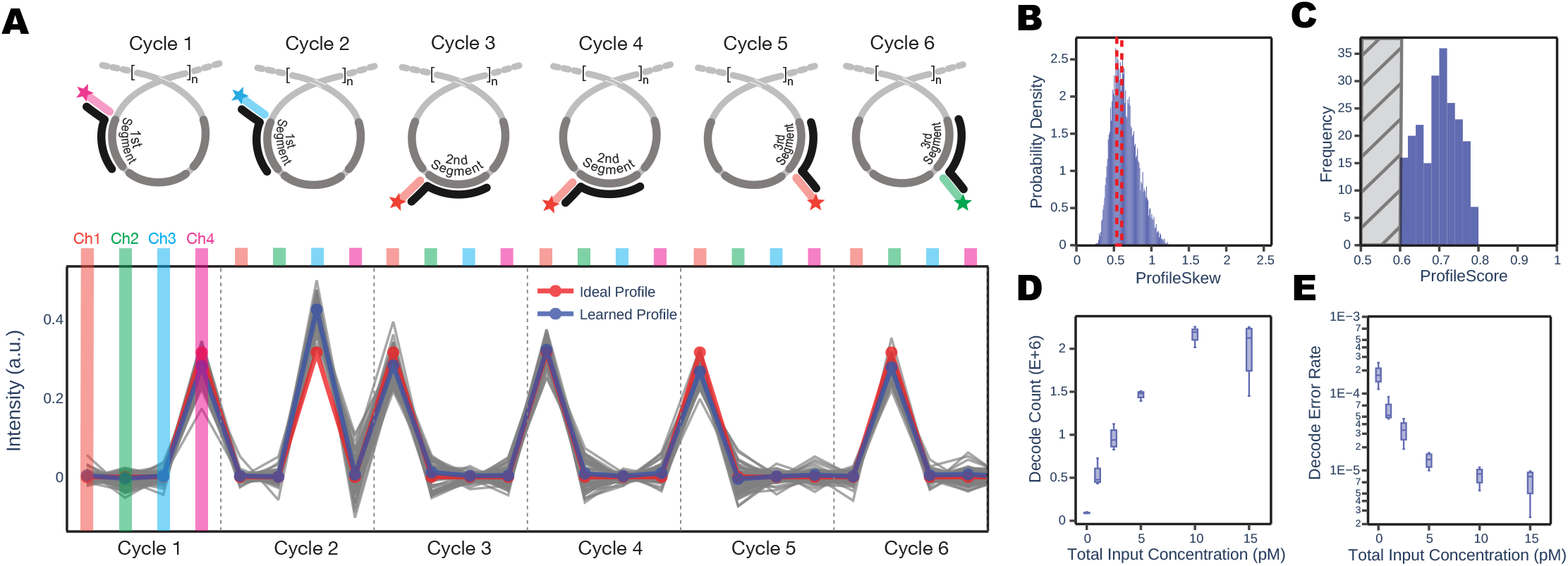
Hypercoding strategy to enable accurate, high-plexity readout of RCPs. A) Cyclic probing of RCPs with detection oligonucleotide complexes. After each of the 6 hybridization and imaging cycles depicted, the fluorescent intensity across channels for identified RCPs is recorded to generate a discrete intensity vector. Representative intensity vectors for 100 RCPs (gray traces) that decode to the same Hypercode (profile score >= 0.7). The learned, consensus Hypercode profile (blue trace) and ideal intensity vector (red trace) are superimposed to reflect the systematic deviations in the measured intensities associated with this Hypercode. B) A probability density function of the distance, or Profile Skew, between ideal and learned Hypercode profiles. A distribution of profile scores for RCPs that decoded to the Hypercode associated with the code 4311122434. RCPs with profile scores below 0.6 are not assigned to a Hypercode to avoid errors from incorrect assignments. C) Histogram of Profile Score, or ratio of the smallest distance to that of the sum of the smallest and second-smallest distances matching valid Hypercode profiles, for all RCPs decoding to the Hypercode associated with the code 4311122434. D) The number of decoded RCPs over a range of target input concentration spanning 0 pM to 15 pM. E) Decode error rate as a function of target input concentration spanning 0 pM to 15 pM.

The number of distinct Hypercode profiles scales with the number of cycles and resolvable optical signatures at each cycle, termed states, thus dramatically expanding the potential assay plexity (Extended Data Table 1). We considered designing Hypercodes with larger expected intensity differences, however this reduces panel plexity (Extended Data Table 2). For this work, we designed Hypercodes within the framework of a 4-state system and varied the number of segments to achieve plexities up to 12,000 Hypercodes^19,20^ (Methods).

### Decoding Performance

To characterize the performance of Hypercodes at different plexities, we constructed two Plenoid libraries: a mid-plex library of 1,048 Hypercodes (1K plex) and a high-plex library of 12,000 Hypercodes (12K plex) (Methods). All Plenoids targeted the same synthetic DNA target to facilitate direct assessments of Hypercode performance.

Accurate decoding relies on unambiguous and consistent classification of intensity vectors to their respective Hypercodes. Noisy and low intensity vectors initially proved challenging to correctly classify (Extended Data Fig. 1). We implemented machine learning in our decoding algorithm to compensate for the systematic signal variations from ideal profiles, termed profile skew, on a per Hypercode basis (Fig. 2b, Methods). Fig. 2c shows the resulting profile score distribution of RCPs decoding to the Hypercode depicted in Fig 2a. The use of both empirical intensity profiles in decoding and an error-correcting Hypercode set contributed to the highly accurate readout of both panels, including the decoding of challenging RCP profiles. We successfully decoded hundreds of thousands to millions of RCPs across a 10-fold range of DNA inputs per field of view (Fig. 2d).

Estimation of decode errors using the fraction of decoded RCPs assigned to Hypercodes reserved for the detection of decoding errors was below 1 in 100,000 for wells with >1M decoded RCPs (Fig. 2e). We also sought to determine the ability of the profile decoder to perform in the event of a missing cycle, which could happen for a variety of reasons. For nearly all the Hypercodes within the Plenoid library, excluding one cycle reduced our ability to correct for discrete errors^21^. The decode error rate and the number of Hypercodes covered for both the 1K and 12K libraries was robust to partial information loss (Extended Data Table 3).

We next evaluated Plenoid library performance on two key metrics: uniformity of coverage across the individual Hypercodes within a well and reproducibility across wells. Overall decode count is consistent from well to well with a coefficient of variation (CV) of 10.8% (n=8) or 7.8% (n=4) for the 1K and 12K plex data, respectively. Within a well, we observed similar numbers of RCPs decoded for different Hypercodes, with a Hypercode-to-Hypercode CV of 9.8% (1K plex) and 28.1% (12K plex) (Table 1). For the 1K plex data, we detected 1,032 of 1,048 Hypercodes (98.5%) with coverage of ≥100. For the 12K plex data, we detect 11,293 of 12,000 Hypercodes (94.1%) with coverage 10 or greater. For both Plenoid libraries, negative controls with no target DNA covered just 3 (1K plex) or 5 (12K plex) Hypercodes.

**Table 1.** Decoding performance metrics for 1K plex and 12K plex Hypercode libraries. Hypercodes are considered detected when observed more than 10 times within a well. Median coverage is reported for detected Hypercodes (codes with coverage >10). Decode count CV represents the overall CV of total counts from well to well. Hypercode-to-Hypercode CV represents the CV of counts across Hypercodes present in the Plenoid library hybridizing to the same target. Normalized Hypercode count CV normalizes for the relative abundance of each Hypercode measured across wells. Decode error rate is the estimation of the probability that any decode call is incorrect (Methods).

| Panel |  | Total Decoded RCPs | Number of Hypercodes with coverage >10 | Median Hypercode counts | Decode count CV (%) | Raw Hypercode count CV (%) | Normalized Hypercode count CV (%) | Decode Error Rate |
| --- | --- | --- | --- | --- | --- | --- | --- | --- |
|  | 12K Plex | 1,036,565 | 11,294 | 86 | 7.8 | 50.9 | 28.1 | <1 in 100,000 |
|  | 1K Plex | 1,959,438 | 1,046 | 1,620 | 10.8 | 79.9 | 9.8 | <1 in 100,000 |

### Genotyping

To evaluate the potential for Hypercoding to be used as a novel genotyping method, we developed a panel of Plenoids consisting of 564 Plenoids targeting 209 small variants (200 SNVs and 9 indels) in 38 genes critical for pharmacogenomic (PGx) testing (Clinical Pharmacogenetics Implementation Consortium, Supplementary Table 1). The panel covered targets in a range of genomic contexts including a range of GC content (33%-57%, ∼40 bases around ligation site), repeat motifs (Extended Data Fig. 2), and loci with high similarity to other regions of the genome (e.g. *CYP2D6* and the pseudogene *CYP2D7*). For standard genomic loci, a Plenoid with a unique Hypercode and allele-specific 3’ nucleotide is designed for each reference and alternate allele and genotypes are assigned based on the relative abundance of counts for each Hypercode (Fig. 3a, Methods).

**Figure 3.**
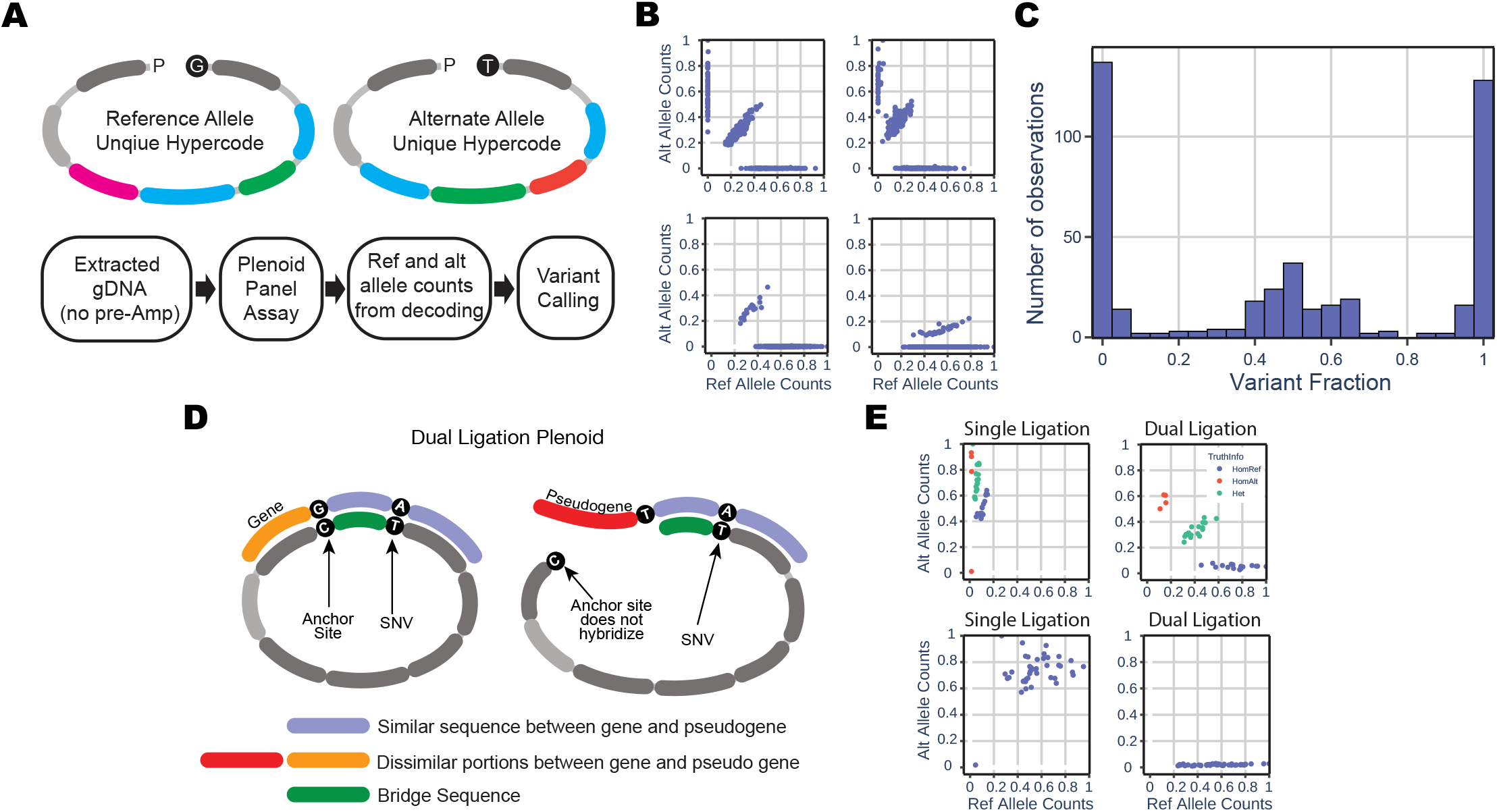
Germline genotyping using Hypercoding technology. A) Description of genotype strategy using Hypercoding technology. Two Plenoids with unique Hypercodes are designed for each variant of interest. The Pleno assay workflow described in Figure 1 is executed using intact non-amplified genomic DNA as input. Decoding process produces counts for the ref. and alt. Plenoid for each of the variants in the panel. Variant fraction calculations and variant calling are performed to produce genotype calls. B) Examples of raw normalized ref. and alt. counts for four variants showing distinct clusters Homref, Het and HomAlt genotypes. Upper left, example of SNV variant (rs4680) with high global minor allele frequency (GMAF) of 0.390. Upper right, example of SNV variant (rs1801265) with GMAF of 0.231. Lower left, example of INS variant (rs41303343) with low GMAF of 0.0293. Lower right, example of DEL variant (rs35742686) with low GMAF of 0.00918. C) Distribution of measured variant fractions across a panel of 213 variants measured across 108 samples showing three well separated populations. 150 measurements from each genotype were downselected to better graphically represent each of the HomRef Het and HomAlt fractions. D) Description of the strategy to genotype variants in regions with high homology (pseudogenes). Two ligations are required to form a circle: one ligation event occurs at the base of interest and a second ligation event happens at a downstream site where the high homology region and the region of interest differ. A bridge oligo is added to fill the gap between the two ligation sites. E) Examples of variants in high homology regions genotyped using standard single ligation vs. double ligation Plenoids. A variant (rs774671100) with low GMAF where known HomRef genotype samples show elevated counts for the alt. allele in the single ligation Plenoid case. Off-target counts are dramatically decreased by using a double ligation Plenoid. A variant (rs3745274) with high GMAF showing all three clusters overlapping and indistinguishable when using single ligation and well separated when using double ligation Plenoid.

Small variant genotyping performance was assessed using a cohort of 108 reference samples from the 1000 Genomes database (Purchased from Coriell, Supplementary Table 2) to determine the performance of small variant calling and to measure detection accuracy. Using 100ng of input DNA, average number of RCPs decoded per sample were 704,515±129,000 with each locus covered by a median of 1,734 RCPs. Hypercoding enabled clear differentiation of 3 distinct clusters corresponding to homozygous ref, heterozygous and homozygous alt samples across SNVs and INDELs as shown on selected examples (Fig. 3b). Variant fraction distribution across the whole panel was consistent showing clear separation among genotypes (Fig. 3c) with average per sample call rates and accuracy of 99.8% and 98.7%, respectively (Extended Data Table 4).

Targets such as *CYP2D6* share highly similar sequences with other genomic loci and present a challenge for NGS and other genotyping methods^22^. To accurately genotype these high homology regions, we developed dual ligation Plenoids that include a 3’ targeting arm specific to a SNV of interest and a 5’ targeting arm that hybridizes to a nearby anchor site (Fig. 3d). Anchor sites are chosen based on low levels of polymorphism and the presence of a single nucleotide that distinguishes between the target of interest and the homologous region (Methods). A bridge oligonucleotide hybridizes to the genomic sequence between the target site and anchor sites, thus requiring two ligation events per Plenoid: one at the target site and one at the anchor site (Fig. 3d). In the case of standard Plenoids, off-target ligation prevents clear cluster separation whereas dual ligation designs significantly improved detection fidelity (Fig. 3e). For two representative loci, readout error rates in homozygous reference samples were reduced from 72% to 5% on average, validating the ability to detect variants from the region of interest in a background of highly homologous sequences without the need to perform upfront disambiguation using multiplex PCR amplification or target enrichment.

### Copy Number Variations

Next, we used Hypercoding to detect whole- and sub-chromosomal copy number variations (CNVs). For sub-chromosomal CNVs, we included 16 Plenoids across 4 different regions of the clinically relevant *CYP2D6* gene in the PGx panel to enable detection of amplification, deletion, and hybrid CNVs (Fig. 4a). Copy number (CN) changes were detected by normalizing observed detection events in *CYP2D6* CNV Plenoids to regions of the genome with stable copy number and comparing to control CN=2 samples (Methods). Figure 4a depicts the measured CNV signal for whole gene deletion (CN=1) and amplifications (CN=3 and 4) across 4 separate gene regions demonstrating good separation across samples. Highly challenging *CYP2D6* hybrid alleles that require sub-gene resolution were also resolved using Hypercoding-based estimation (Fig. 4b). The Hypercoding-based called CN values were 99.8% concordant with the expected CN as determined by NGS (Extended Data Table 5).

**Figure 4.**
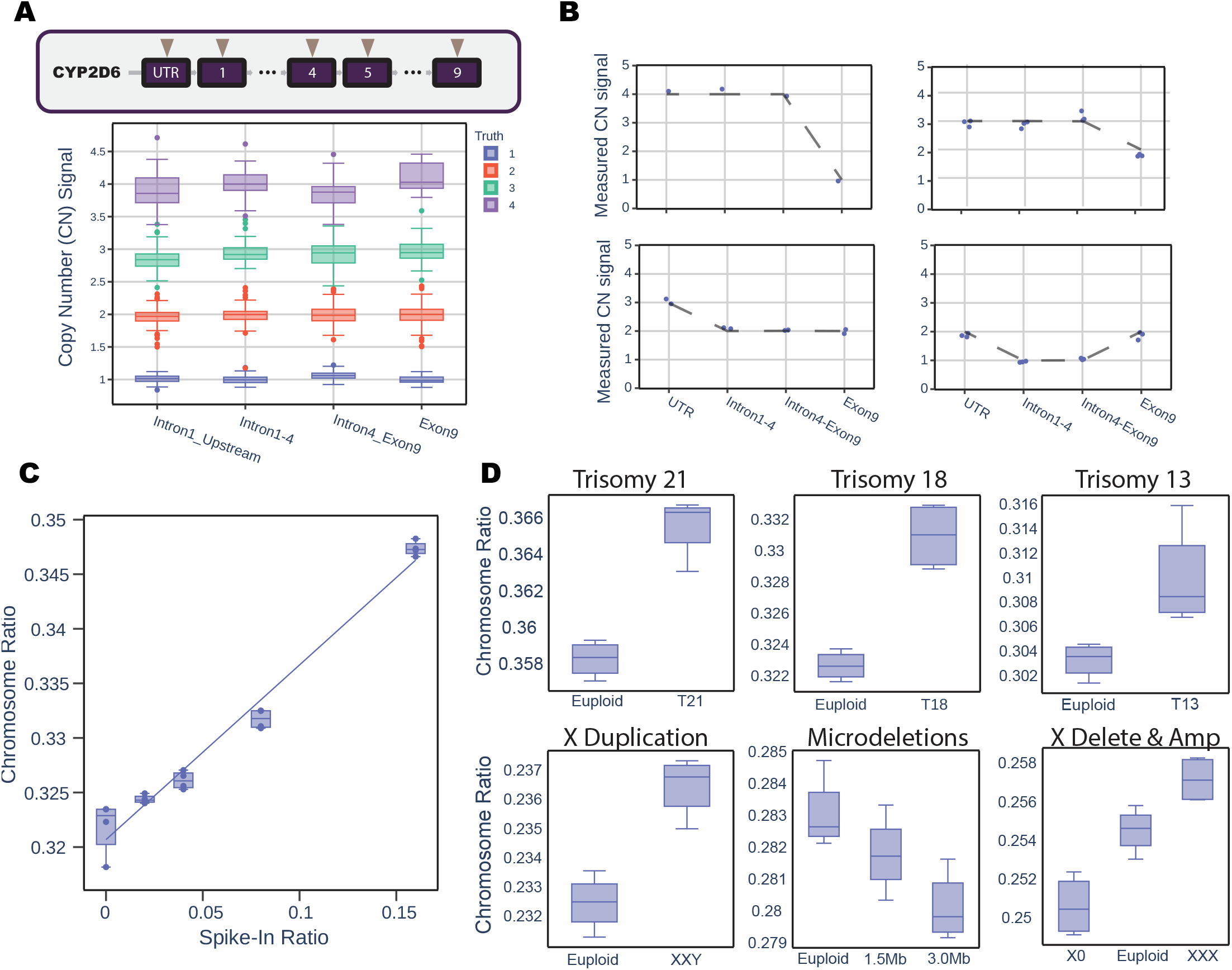
Copy number variation determined using Hypercoding. A) Whole gene copy number variations (CNVs) detection in clinically relevant highly homologous CYP2D6 gene. Locations within CYP2D6 where Plenoids were designed are indicated with arrows. Measured copy number across regions of interest of CYP2D6 shows distinct clusters of whole gene deletions (CN=1) and whole gene amplifications (CN=3 and CN=4) (n=108). Color represents known truth using NGS. B) Detection of CYP2D6 hybrid alleles show partial deletions or partial amplifications of different regions within the gene demonstrating sub-gene resolution capabilities of Hypercoding. Upper left, Exon 9 conversion *10+*36/*36+*36 for sample NA18757; upper right, Exon 9 conversion *10+*36 for samples NA18553, NA18624, NA18612; lower left, intron1 *68 for sample NA12878; and lower right, *68 and exon9 hybrid *13 for sample NA19982. C) Aneuploidy detection using chromosome ratio in contrived samples for NIPT applications. Fetal fraction of 0 – 16% for Trisomy 21 spike-ins with R2 = 0.969. D) Aneuploidy detection of different genetic conditions at 4% spike of fetal DNA. Sub panels, from left to right, top to bottom: Euploid vs 4% spike of Trisomy 21, Trisomy 18, Trisomy 13, X duplication, large and small microdeletions, and X deletion and amplification.

To determine whole-chromosome CNV capabilities and to assess the ability to detect small quantitative changes, a Plenoid panel targeting common aneuploidies relevant for non-invasive prenatal testing (NIPT) was designed by tiling multiple Plenoids across several chromosomes of interest. For validation, contrived samples at a total input of 15ng were developed mimicking different levels of fetal fraction by spiking varying amounts of an aneuploid sample into the background of a euploid sample. We detected aneuploidies from euploid background titrating down from 15% down to 2% fetal fraction for trisomy 21 (Fig. 4c). In all contrived samples for trisomy 13, trisomy 18, trisomy 21, XXY syndrome, chromosome 22 large and small deletions, monosomy X, and trisomy X, there is a difference between the euploid sample and the 4% mock fetal fraction sample (Fig. 4d), which demonstrates the capabilities of Hypercoding to detect whole chromosome CNVs.

### Target Quantitation

To demonstrate the platform’s ability to quantify sample target concentration, a 21-plex Plenoid panel was constructed to synthetic targets. A calibration curve spanning 1,400-fold dynamic range from 25 fM to 35 pM for each target exhibited strong linear correlations (R^2^ > 0.99, Extended Data Fig. 3) with reproducible slopes (average CV of < 4%, n = 6). The lower and upper limits of detection (LoD) of the panel were determined to be 25 fM and 100 pM respectively, with more than half of individual targets detectable <10 fM using a previously described definition of the lower LoD^23^ (Fig. 5a, Extended Data Table 6).

**Figure 5.**
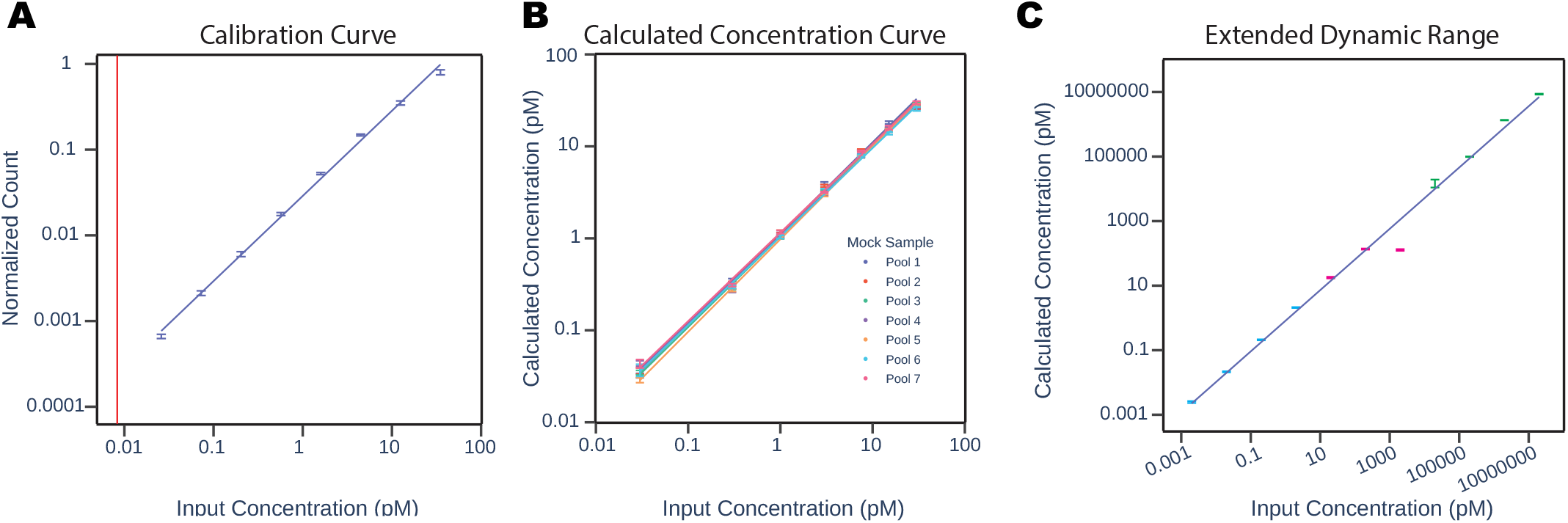
Demonstration of target quantification using Hypercoding. A) Normalized RPC counts across 3 log of target input (0.03-30 pM) demonstrate a linear response (R2=0.997) of decoded RCPs when using internal calibrators to compensate for well-to-well variability and the effects of surface saturation at high target input concentration (n=21). All targets are above the system limit of detection (meanBlank + 1.645*STD_Blank_ + 1.645*STD_lowest concentration_). B) Correlation plots of the calculated concentration for each of 21 targets vs. target input concentration for seven contrived samples. Each target independently exhibits high concordance (R2 > 0.99). Concentrations were calculated via comparison of normalized target counts to a calibration curve. C) Demonstration of extended dynamic range quantitation of 11 targets spanning 10 log (1 fM – 10 μM) within a single sample utilizing one, two, or three separate wells to perform sample serial dilutions.

Targets were quantified by designing 7 contrived samples containing all targets at varying concentrations spanning 30 fM to 30 pM (Methods). Robust linear correlations (R^2^ > 0.99) were achieved when comparing the input target concentration within the sample to its calculated concentration (Fig. 5b). Error of quantification for calculated concentration ranged from 42.5 ± 44.2% (30 fM, all targets) to 13.1 ± 4.6% (30 pM, all targets) across the entire 3 log dynamic range. Inter-assay (plate-to-plate) variability of calculated concentrations for targets was also low, with CVs across all targets ranging from 25% (30 fM) to 12% (30 pM) (Extended Data Table 7).

We further challenged the dynamic range capabilities of the platform by serial dilution of a sample with targets that span 10 logs in input concentration. This sample and its two dilutions were then assessed over 3 wells such that each well spans a particular 3-4 log range of the extended dynamic range (Methods). The resulting calculated concentrations across the entire dynamic range were highly concordant with input concentrations (R^2^ > 0.98) demonstrating the ability to the platform to detect targets spanning a concentration range of 10 logs (Fig. 5c).

## Discussion

We present Hypercoding, a technology for detecting and quantifying thousands of DNA analytes in parallel in a single well without the need for target enrichment or cumbersome library preparation. Hypercoding readout is rapid, inexpensive, and readily enables for the readout of thousands of targets in parallel while producing genotyping and quantitation results that match truth sets generated using digital PCR^24,25^ and NGS.^26^

The Hypercoding system was implemented on Pleno’s proprietary benchtop instrument (RAPTOR™), which integrates fluidics and imaging modules with a long-range X-Y-Z motion subsystem and computational engine and implements the described readout chemistry workflow. This combined detection and decoding approach enables the Hypercoding platform to have very rapid readout cycles while maintaining robust data quality. Samples are loaded into a multi-well plate making the upfront assay compatible with existing automation tools, which simplifies deployment in working laboratories.

Leveraging a simple hybridization-based detection workflow eliminates the need for multi-component reagents and enzymatic chemistries^27,28^ by using a small number of detection oligos. Detection schemes can be tuned to address assay requirements such as minimized decode error rate, reduced readout time, higher plexity in the tens of thousands, and/or improved dynamic range quantification of a small number of targets by varying the number of fluorescence states and cycles. Additionally, the novelty of machine learning-based Hypercoding allows for systematic noise compensation and robust assay readout error correction while maintaining low decoding time for high plex assays.

Universal Plenoid design principles can be applied to a wide variety of content of interest, which enables customization to meet the needs of diverse applications. Genomic coordinates are readily turned into a Plenoid panel design, which can be commercially synthesized rapidly and economically for targeted testing. In contrast, similar technologies like microarrays require significant development efforts to alter the content, customize probe designs, and manufacture the final substrate. Rapid design-to-test turnaround drastically speeds up assay development cycles, increasing the pace of scientific discovery and commercialization.

In this work, we demonstrate accurate genotyping and copy-number calling of PGx-relevant genes to illustrate the utility of Hypercoding for omics applications. The use of dual ligation Plenoids helps overcome the limitations of standard microarray assays for querying targets with homologs such as the clinically important gene *CYP26D*^24,25^. Further integration into commercial interpretation and reporting software is possible from the generated industry standard variant caller output file formats (e.g., .vcf). Taken together, the combination of a streamlined wet-lab workflow, a purpose-build readout instrument, and processing algorithms enables the production of clinical reports from extracted genomic DNA in under 8 hours.

Hypercoding is highly quantitative and precise for both relative and absolute measurements given that each detected RCP originates from a unique Plenoid-target interaction, and thus a single copy of the target in the original sample. This technique provides two advantages over dPCR. First, the classification of thousands of discrete RCP types far exceeds the multiplexing ability of existing dPCR^8^. Second, the simultaneous detection of millions of individual RCPs overcomes the traditional limitations of molecular partitioning and sample dilutions required for dPCR. Newer dPCR methods have overcome some of these limitations and achieved 6 logs of dynamic range, though at limited target plexities^11^.

Furthermore, Hypercoding can be used for quantification of proxy analytes, as is now becoming common in proteomics applications^29^. We demonstrated the detection and accurate absolute quantification for 21 proxy-analytes over concentrations spanning 3 orders of magnitude and dynamic range extensibility over 10 logs using multiple wells, which covers the range of commonly tested blood proteins^30^. In addition, the use of internal calibrators to compensate for well-to-well variability and mitigate the effects of surface saturation at high target input concentration further extends the linear dynamic range of the platform with minimal impact to system output.

In this study, we utilize the digital counting advantage of Hypercoding to estimate copy number variation and achieve accurate quantification of whole-gene and sub-gene amplifications and deletions, including whole chromosome aneuploidy. Furthermore, we show good performance in challenging conditions such as genomic regions with high homology and high GC content (*CYP2D6*) as well as low target abundance down to 4% simulated fetal fraction. To achieve detection at even lower abundance, assay variability can be further reduced by averaging the signal from separate Hypercodes targeting multiple sites within a region of interest. Hypercoding as presented in this manuscript has several limitations. First, the estimation of decode errors in the panel is a measure of decoder performance and does not include biochemistry driven errors such as mis-ligations and amplification challenges, both of which are limited by the performance of commercially available enzymes^31^. Second, thought must be given to the genomic proximity of targets within a panel; we have observed interference from close proximity (<20 nucleotides) variants negatively impacting the performance of one or both targets of interest. Third, due to limitations in long (>100-mer) oligo synthesis, an entire Plenoid pool often includes truncated products that cannot participate in circularization. The prevailing strategy for solving the noted challenges is through design iterations, dual ligation designs, alternative Hypercode assignment, and oligo purification. Future developments of Hypercoding as a technology on the RAPTOR platform are focused on addressing these critical to quality aspects.

Further technology development will include extensions of the system to additional analyte types including methylation states, RNA, proteins, and indirect readout of multi-omic targets. Detection of more analytes per sample using higher-plexity panel can be achieved by increasing the number of fluorescent colors (or states), a combination of colors, or color amplitude modulation. While the current detection and decoding pipeline can detect and decode ≥2M RCPs/well, scaling to >10M is possible through image tiling, improved optics, and custom well plate geometries.

In summary, Hypercoding is an innovative technology for detecting and quantifying thousands of DNA targets simultaneously within a single well without the need for target enrichment or complex library preparation. The Hypercoding technology offers an alternative to traditional NGS or dPCR technologies in the diagnostics space. Offering scalability, low cost, and automation potential, Hypercoding provides a powerful platform for clinical, research, and commercial applications.

## Author Information

These authors contributed equally: Madhavi Bathina, Angela P. Blum, Jeff Brodin, Nicole DeBuono, Samantha Snow, Bo Lu, Matthew R. Naticchia, Daniel Ortiz, Andrew Richards, Cyrus de Rozières, Rachel Schowalter, Sarah Shultzaberger, Stephen Tanner, Christy Trejo, Ludovic G. Vincent, and Spencer Ward.

## Contributions

The author list is divided into four sections, the first three of which are in alphabetical order:

Category 1: Authors in this group contributed to the core technological development and the writing of the paper.

Category 2: Authors in this group made specific contributions to certain parts of the paper.

Category 3: Authors in this group contributed to the development of the underlying instrumentation that enabled the studies performed in this paper, provided operational support, or significant editorial help.

Category 4: Authors in this group provided supervision and/or foundational concepts for the studies.

## Online Methods

### Sample Preparation for Readout

#### Overview

Preparing samples, including converting linear Plenoids™ to circularized Plenoids in the presence of a target sequence and amplification to rolling circle products (RCPs), consists of three steps: ligation, exonuclease digestion, and rolling circle amplification (RCA, each described below). The ligation and exonuclease digestion steps are performed sequentially in the same well of a polymerase chain reaction (PCR) plate. The sample is then transferred to a custom 96-well plate with an optically clear bottom and a DNA-binding surface for sample capture followed by RCA.

#### Ligation

The DNA sample to be probed was combined with a Plenoid pool, a buffer solution and a high-fidelity thermostable DNA ligase in a 20 μL reaction volume (10ng - 100ng of DNA). Mixtures were cycled between 95 °C and 60 °C three to six times with 2-minute incubations at 95 °C to denature duplex DNA and 20-minute incubations at 60 °C to allow for target hybridization and ligation.

#### Exonuclease Digestion

Each 20 μL ligation reaction was diluted with 30 μL of an exonuclease mixture to specifically digest only linear DNA. The ligation and exonuclease mixtures were then incubated at 37 °C for 30 minutes followed by 95 °C for 5 minutes to inactivate the enzymes.

#### RCA

Prior to amplification, 50 μL of the exonuclease-treated ligation sample was transferred to a different well of a polymer-treated glass-bottom plate and the plate was incubated at 42 °C for 60 minutes to allow for surface binding of the circles. Next, the solution in each well was removed and discarded prior to the addition of 50 μL of RCA reaction mixture, which contained a sequence-specific primer, dNTPs, a buffer solution, and Phi29 DNA polymerase. The plates were then incubated at 42 °C for 1-2 hours. After this period, the reaction was quenched by washing with a Tris-EDTA solution.

### Readout Instrument

Readout was done on the RAPTOR™ instrument, a Pleno-designed integrated opto-mechanical and fluidics instrument. The fluidics subsystem integrates an eight-channel plate washer, an eight-channel strip buffer dispense module, ten independent precision dispensers, and a waste-management system. The optical subsystem is a four-color imager with two excitation paths at 532 and 639 nm and two cameras for capturing images of two spectrally separated wavelengths per excitation path.

To prepare the plate for detection, any remaining Tris-EDTA solution from the RCA step was aspirated from all wells of the Decode Plate. Stripping solution (200 μL, alkaline solution, pH>12) was added to each well and incubated for 60 seconds. Wash buffer (150 μL), an alkaline buffered solution with monovalent salt, EDTA and surfactant, was added to each well to dilute the stripping solution followed by aspiration of the stripping and wash buffer mixture. All steps within the process are done at ambient temperature within the instrument.

1) Hybridization was initiated by dispensing readout cycle reagents (50 μL) containing 16 detection oligos in a buffered solution (containing mono- and di-valent salt, EDTA, and surfactant) into each well. The plate was incubated for 9 minutes to allow for hybridization of detection oligos to the RCPs. The readout reagents were aspirated and then wells were washed 10 times with wash buffer (260 μL) and left in wash buffer for imaging (Fig 1b). 2) The plate was moved over the imaging module, with the camera centered on the first well to be imaged. For each detection wavelength a fluorescent image of the RCPs was captured, and the process repeated for all wells. 3) Following imaging, the wash buffer was aspirated, and the wells were incubated with stripping solution (200 μL) for 60 seconds. The stripping solution was diluted with wash buffer (150 μL), aspirated, and then the next readout cycle reagent was added. Steps 1-3 were repeated for eight to ten cycles depending on the code length, with unique detection oligos in each readout cycle.

### Codespace Design

Given *S* symbols (corresponding to distinct system fluorescence states) and *c* readout cycles, there are *S*^*c*^ possible *codewords*. A strength of decoding by hybridization is that the number of symbols *S* can be increased by using more types of fluorophore molecules, combinations of fluorophores, or fluorescence levels. Furthermore, the number of cycles can readily be decreased for faster readout or increased for higher target plexity. The one-hot encoding scheme, where each RCP fluoresces in exactly one of several colors, was selected for ease of high-confidence decoding.

The list of possible codewords is filtered using heuristics based on the methods used in the readout assay. For example, we exclude low complexity codewords that contain long runs of a single symbol out of concern that such codewords are more vulnerable to being misread through biochemistry or optical artifacts.

Given the collection of valid codewords, we now select a *codespace* – a collection of codewords satisfying the minimum pairwise Hamming distance threshold (HD^min^ = 3 for this work) to enable error correction^1^. Selecting a maximum-size codespace, which enables the largest possible assay plexity, is difficult since enumerating all code combinations becomes computationally intractable for nontrivial cases. There are theoretical upper bounds on how large such a space can be^2^, but these bounds do not assist with constructing a maximum-size codespace. In practice, the maximum achievable codespace is often smaller than the known theoretical upper bound. A practical approach is to heuristically generate multiple codespaces, then select the candidate codespace with the largest number of codewords. One approach to design a codespace is to begin with an empty list of selected codewords, and an available list of all valid codewords, generated as above. We select a codeword at random from the list of available codewords and add it to the selected codewords list. We then remove from the list of available codewords any candidates whose Hamming distance from the chosen codeword is less than a chosen cutoff. Random codeword selection and subsequent pruning of the available codewords list is repeated until there are no more available codewords. This entire codespace generation process can be repeated numerous times to generate codespaces consisting of different sets of codewords. Modifications to the random selection criteria have been proposed to obtain larger codespaces^3^.

Our implementation of the codespace generation procedure evaluates 10,000 candidate S=4 and c=8 codespaces in 3.7 hours on a 3.6 GHz desktop PC, with the maximum codespace size obtained by the 4721^st^ iteration after 1.75 hours. The final maximum codespace was a total 1167 codewords (1K Plex). A similar exercise was repeated for S=4 and c=10, and yielded a total of 13,156 codewords (12K Plex). Runtime increases for still larger symbol or cycle counts but can readily be accelerated by using additional threads.

### Hypercode™ Design

Each Hypercode is a nucleotide sequence made up of several short oligos, or segments, which are used to identify a Plenoid through hybridization with fluorescently labeled detection oligos. The set of all these segments, which collectively encode all the codewords of the codespace, is referred to as the *segment set*. The design of a highly sensitive and specific segment set requires careful selection of sequences with favorable biochemical properties. Each segment is screened for robust hybridization to its complimentary detection oligo while exhibiting low affinity to probes, to detection oligos for other segments, and to other segments within the segment set.

Given segments of length *L* in a Plenoid, there are 4^*L*^ possible nucleotide sequences to form segments, which is a very large set for oligonucleotides *L*>12. From an initial set of oligomers with *L*=18 (∼68 billion sequences), we disallow sequences that are anticipated to be vulnerable to synthesis errors (e.g. homopolymers, high GC content, complex secondary structures). Plenoids with long stretches of dinucleotide and trinucleotide repeats are excluded as well. Next, we impose a Hamming distance cutoff between segments to ensure sequence diversity and strictly reduce the segment set to a more manageable set (∼10^5^-10^6^ sequences). Segments are then screened for homology against the human genome and near matches removed. The final segment set included over a hundred candidate sequences. For this work, 16 sequences from the final segment set were assigned to a Hypercode position for a total of 80 sequences across 5 positions.

For the 12,000 plex library, Hypercodes were then constructed by concatenating 5 segments, one from each of the 5 sets of 16 segments, to match a codeword from the codespace described above. For instance, codeword 43-12-34-41-23 was assigned segment #15 from the segment set corresponding to position 1 as well as segments #2, #12, #13, #7 from segments sets 2-5.

### Plenoid Design

The design workflow for a set of Plenoids for a particular panel starts by choosing a list of loci within a species of interest. These loci can be read from a standard VCF file^6^ that contains the chromosome, position, locus id, reference allele, and alternate alleles. Generally, each locus is probed by N probes where N is the number of alleles (reference and alternate).

For each potential probe, we query the reference genome to obtain the target sequence. The target sequence is then divided at the locus of interest into 2 separate sequences: the sequence that is toward the 5’ end is considered the 5’ target of the locus and the sequence towards the 3’ end is considered the 3’ target. For the 5’ probe/target pair and the 3’ probe/target pair, we select the pairs that have a T_m_ slightly above the ligase operating temperature but below the denaturation temperature. The selected reverse complement sequence for the 5’ target is considered the 5’ probe arm, and similarly the 3’ probe arm for the reverse complement of the selected 3’ target. At this point each target has 5’ and 3’ probe arms that are ready to be assigned to a Hypercode. To minimize potential intramolecular interactions between the probe arms and Hypercode, a series of potential assignments must be computed and screened for secondary structure. The combination of a Hypercode and a pair of 5’ and 3’ probe arms form a linear Plenoid.

Consideration must also be given to pseudogenes or other regions of the genome that have high sequence similarity with the target locus and polymorphisms across populations. Due to the high similarity, a standard single ligation Plenoid is likely to hybridize on the similar sequence, undergo ligation, and result in off-target counts being generated. To fix these issues of sequence similarity, we utilize the Dual-Ligation Plenoid designs that can distinguish the similar sequences from the target of interest by using an additional oligo, which we call a bridge, to be used with a Plenoid (Fig. 3d).

### Image Analysis and Signal Processing

The imaging of fluorescently tagged detection oligos bound to the RCPs is performed on the RAPTOR platform. The decoding process begins with analyzing these images across readout cycles and channels. Subsequent analysis steps involve intensity normalizations, feature identification, intensity extractions, and data conditioning operations. Minimizing optical, electrical, biochemical, thermal and motion-induced noise sources is crucial for optimizing decoding performance. To this end, the processing pipeline employs subpixel feature detection methods, akin to those described elsewhere^8,9^. Subsequently, subpixel registration techniques are applied to pinpoint the same RCP in all readout cycles and channels^10,11^. Optical and biochemical imperfections, such as nonlinear distortion and non-uniform illumination, are corrected for in-situ or via prior calibrated correction factors. Finally, the extraction of intensities is done by convolving the feature image raw pixels with an appropriately tuned extraction kernel. These computationally intensive processing steps are implemented using C++ and Halide^12^ to enable pseudo-real-time processing.

Following extraction, the raw intensity values from the RCP-population undergo normalization. We start by mapping all intensities into a standard range to eliminate outliers, for example into the [0.5, 99.5] % range of all intensity values. Next, we correct for spatial variability in foreground and background intensity resulting from factors such as non-uniform illumination. The image is divided into a grid of sub-images, and high and low percentiles of intensity in each subregion are measured. RCP intensities are re-scaled such that, after correction, the background and foreground intensities of the various subregions are comparable. To minimize edge effects across subregions, bilinear interpolation is applied to the correction factors. The use of color-balanced control Plenoids can help ensure that all fluorescent color channels observe some bright objects in each readout cycle, regardless of the true application sample plexity and the sample assayed. In a third step, color crosstalk caused by the overlap of emission spectra of different fluorophores is corrected^13,14^.

Color crosstalk can be modelled as a linear mixing operation and correcting for it by “unmixing” the observed intensities^13,14^. These linear models can readily be measured for a particular combination of dye molecules, excitation lasers, and filters used. Once the crosstalk between colors is quantified, it can be corrected by multiplying the received intensities by the inverse of the 4 × 4 matrix of color crosstalk coefficients. Finally, the intensity vector of each RCP is normalized to unit norm.

### Approximate Profile Decoding

#### Error-Correcting Decoding

A simple decoding method is to determine, for each readout cycle, the most likely symbol or (fluorophore state). If the sequence of symbols **Q** = *q*_*1*_, *q*_2_,… *q*_*c*_ and *q*_*c*_ ∈ *S* perfectly matches a codeword *h*, the RCP is considered decoded with no mismatch. Given ***Q*** we can compute the ideal profile for *h*: a unit norm vector in intensity space, **Q**_ideal_ = *q*′_*1*_, *q*′_2_,… *q*′_*j*_ and j ∈ *S*∗ *c*, where *S* is the number of symbols and *c* the number of cycles. Due to readout noise sources such as non-uniform illumination, or cross-hybridization, there may be errors in the symbols read out for particular RCPs and readout cycles. Traditional error correction enables correction of up to *x* symbol errors in a codespace with minimum pairwise Hamming distance *2x+1*, where we correct the sequence of symbols to the unique valid codeword where at most *x* symbols disagree with the correct symbol set.

This symbol-by-symbol “hard” decoding has limited accuracy. Using knowledge about the probabilistic distribution of the intensities we can apply a more effective assignment and error correction. A probabilistic “soft” decoding assigns codes as

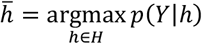

where Y = [*y*_*1*_, *y*_2_,. . *y*_*j*_] and *j* ∈ *S*∗ *c* is the entire *S* colors × *c* cycle *intensity vector* of an RCP and ***H*** is all possible codes in the codespace. This formulation has the advantage that it can probabilistically weigh the effect of different noise sources and track statistical dependencies of intensities across readout cycles. For example, due to secondary structures that might form, a particular Hypercode may exhibit brighter-than-average intensity in certain readout cycles and dimmer-than-average intensity in others.

### Hypercode Intensity Profiles

Even after conditioning the received intensity vector into its normalized form, there still exist signal variations in the sense that even without noise, the intensity of an RCP is not a clean 0/1 sequence of intensity measurements. Instead, RCPs of a given codeword *h* share a common baseline readout intensity vector 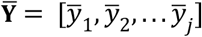 and *j* ∈ *S*∗ *c*, which we call the *Learned Hypercode Profile*. To find the Hypercode Profile for RCP species *h*, we average the unit-norm-scaled intensity vectors of all RCPs that are confidently assigned to codeword *h* using a basic “hard” decoding step. Thus, Hypercode Profiles are derived using empirical intensity data, applying one or more iterations of a clustering algorithm.

We then use intensity profiles to decode RCPs. With evidence that residual noise and errors in the intensities are distributed according to independent Laplacian probability distributions, a good approximation to the probability formula above is:

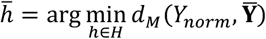

Where *d*_*M*_(*A, B*) is the *Manhattan distance* between two vectors **A** and **B**, defined as

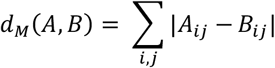

and Y_norm_ is the normalized intensity footprint of a given RCP, and indices *i,j* refer to the Hypercode *i* at intensity profile dimension *j* ∈ *s*∗ *c*. This distance metric is also useful for assessing the confidence in each assignment *–* the lower the distance, the better. We define the *profile score* for a decoded RCP as the quantity 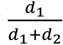, where *d*_1_ and *d*_2_ are the smallest and second-smallest distances to Hypercode profiles. This profile score ranges from a minimum of 0.5 (lowest confidence) to a maximum of 1. This distance metric is useful for characterizing the fluorescence behavior of a given Hypercode: For each Hypercode *h* we compute the distance, *d*_*M*_ between its empirical intensity profile 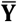 and the ideal intensity profile **Q**_*ideal*_ for its codeword *h* and refer to this distance as the *profile skew* for the Hypercode.

To approach the problem stochastically we employed a matrix-variate Gaussian mixture model to capture the correlation of the fluorescence signals across color channels and cycles, and use this model in a Bayesian estimator called “PosTCode”^15^. Specifically, a prior distribution of the probabilities of targets is assumed. The conditional probability of a given fluorescent signature over all channels and readout cycles given a particular code in the codespace is then modeled as a Gaussian mixture with correlations over the channels and over the cycles. The posterior probability of each code is then computed from the conditional probabilities and the prior distribution. This process is iterated since the correlation matrices in the model need to be estimated also. The model is complex, but typically very powerful if the probability distributions involved are members of the exponential function family. Unfortunately, this model requires fitting many parameters, particularly as plexity increases, and empirical data is not always a good match to this Gaussian mixture model. On the data in this manuscript, we obtained more robust decode performance with the Manhattan distance, rather than the sum-squared distance, which would be appropriate for underlying Gaussian statistics.

### Decode Accuracy and Confidence

To assess accuracy and performance of the integrated Pleno system (chemistry, instrument, and decoding steps), we developed metrics such as estimated error rate and predicted decoding confidence scores. To estimate the error rate, we first measure the **unused error rate**, defined as:

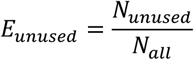

where *N*_unused_ is the number of RCPs assigned to Hypercodes that are known to be absent in the panel, and *N*_all_ is the total number of decoded RCPs for the well. Typically, at least 10% of the codespace is reserved as “unused” space to measure decode error. From the unused error rate, we can estimate the **decode error rate**, the probability that any decode call is incorrect:

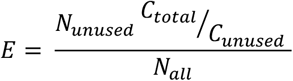

Here we assume that decode errors are distributed uniformly around codewords from the codespace, so the number of decode errors for all codes can be estimated by the number of unused codeword counts *N*_*unused*_ scaled by the fraction of unused codewords in the codespace 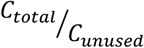.

To ensure high decode accuracy even in the face of readout noise or imperfect spot finding, we filter low-confidence decode calls. We observed that decode errors are correlated with lower absolute RCP intensity, with higher minimum Manhattan distance *d*_*M*_, and with lower profile score as defined above. We used these features, along with labeling of mis-decoded RCPs which correspond to unused codewords, to train a logistic regression model that assigns an error probability to each decoded RCP. We chose cutoffs such that low-confidence RCPs are left un-decoded, yielding high overall decode accuracy (typical decode error rate < 10^-5^) at modest cost to total throughput.

### Decoding Strategies

To demonstrate the effectiveness of error correction, we also truncated the codewords by one symbol producing a codespace with minimum Hamming distance 2, and decoded the nine corresponding rounds of images. Decoding with lower hamming distance resulted in less objects decoded for both the 1K Plex and 12K Plex runs (Extended Data Table 3).

### Germline Genotyping

Germline genotypes were estimated by calculating the variant fraction for each allele in the panel. For this, at least two Plenoids were designed for each variant: one for the reference and one for each alternative allele (a small number of loci contained more than one variant of interest, requiring one Plenoid per variant allele). Variant fraction (VF) was estimated from decoded counts for each Hypercode as:

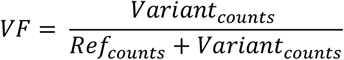

Germline VFs are expected to be 0 for homozygous reference loci, 0.5 for heterozygous loci and 1 for homozygous variant loci. However, in practice each Plenoid exhibits unique performance due to variability in binding and read-out efficiency, and thus calculated VFs might deviate from the expected values. To overcome this, relative performance of reference and alternative Plenoids from known samples with different genotypes at each locus are learned. This information is then used to inform the genotype caller in determining the most likely genotypes for each locus.

### Copy number variation calling

For CNV probes, several normalization steps are performed to detect copy number signals. First, within sample normalization is performed by dividing RCP counts of CNV probes by reference loci in the panel to account for differences in total RCP counts across wells. Second, between sample normalization is performed by comparing normalized counts of CNV probes in test samples with normalized counts in other samples within the same plate. Normalization values can come from either a panel of samples with known copy number (CN =2) across the gene of interest or the median of the normalized counts across plates (assumes most samples in plate are CN=2).

In this study a panel of reference samples was included on each plate for normalization.

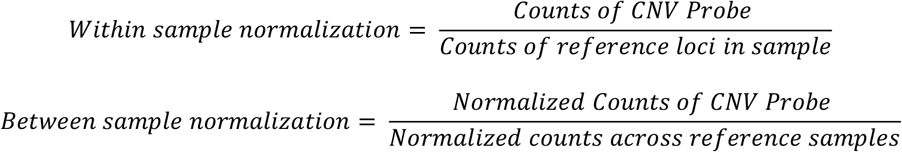

Fold change is calculated for each probe, and the absolute copy number of each CNV region is determined using a Gaussian Mixture Model.

### Star allele calling

Star alleles are functional haplotypes in genes important for pharmacogenomic testing and may involve single variants, combinations of variants, and CNVs. Star allele databases were downloaded from PharmVar. For *CYP2D6*, CNV calling data was combined with small variant data to identify the diplotype of star alleles. For other genes without CNV information, a two-copy state was assumed.

### Target Quantitation

Targets were designed to represent the product of a PCR reaction, wherein a single strand of each duplex would be interrogated by a Plenoid. Each duplexed target was 78 nucleotides in length and featured a region with 30 unique nucleotides for binding Plenoids that is flanked by two common sequences that can be used for population wide amplification or capture. A single Plenoid was designed to hybridize to the unique region of one strand of each target duplex.

To enable quantification of each target, calibration curves were constructed via serial dilution of an equimolar mixture of all 21 target duplexes. The calibration curve for each marker can be linear or sigmoid with the best-fit model selected automatically. For test samples, normalized counts were fitted to the calibration curve for each marker, and the concentration was back-calculated from the calibration curve.

Four control Plenoids were included in all wells (sample and calibration curve) at a constant concentration. These controls served the dual purpose of facilitating optical registration in wells with few RCPs while also enabling normalization of counts for a given target relative to a feature at constant input. The latter enabled extension of the upper limits of detection to concentrations beyond what can physically pack onto the surface of the well plate.

Seven target pools were assembled to assess the quantitative capabilities of the system. In the first seven pools, the 21 targets were divided evenly into seven concentration groups over a three-log dynamic range, spanning 0.3 fM to 30 pM. Across the seven pools, each target was assigned to each concentration group to assess whether all targets could be accurately quantified, irrespective of their concentration. Counts for each target were normalized to the internal controls and concentrations were calculated for each target by comparison to the linear fit of their respective calibration curves.

To assess the target specificity of the Plenoid probes, several drop out control pools were also prepared, wherein all or some targets were deliberately excluded from the pools. Calculated concentrations of the targets present in these pools were highly concordant with input concentrations (R^2^ > 0.98, Extended Data Figure 3) suggesting that the presence of unpartnered Plenoids do not impact quantification. Targets absent from the pool were largely not found, but for those that did register RCPs, their respective calculated concentrations were far below that of the lowest point of the calibration curve (below the defined detection limit).

For the multi-well demonstration, a single sample was prepared representing a wider dynamic range of 10-log, where the 11 targets were distributed evenly into three concentration groupings, spanning from 1 fM to 10 μM. This sample was then serially diluted and assayed in three separated wells. To minimize the number of wells required for quantification, three-point calibration curves were constructed for each target at the relevant concentration range and dilution.

---

**Extended Data Table 1.**
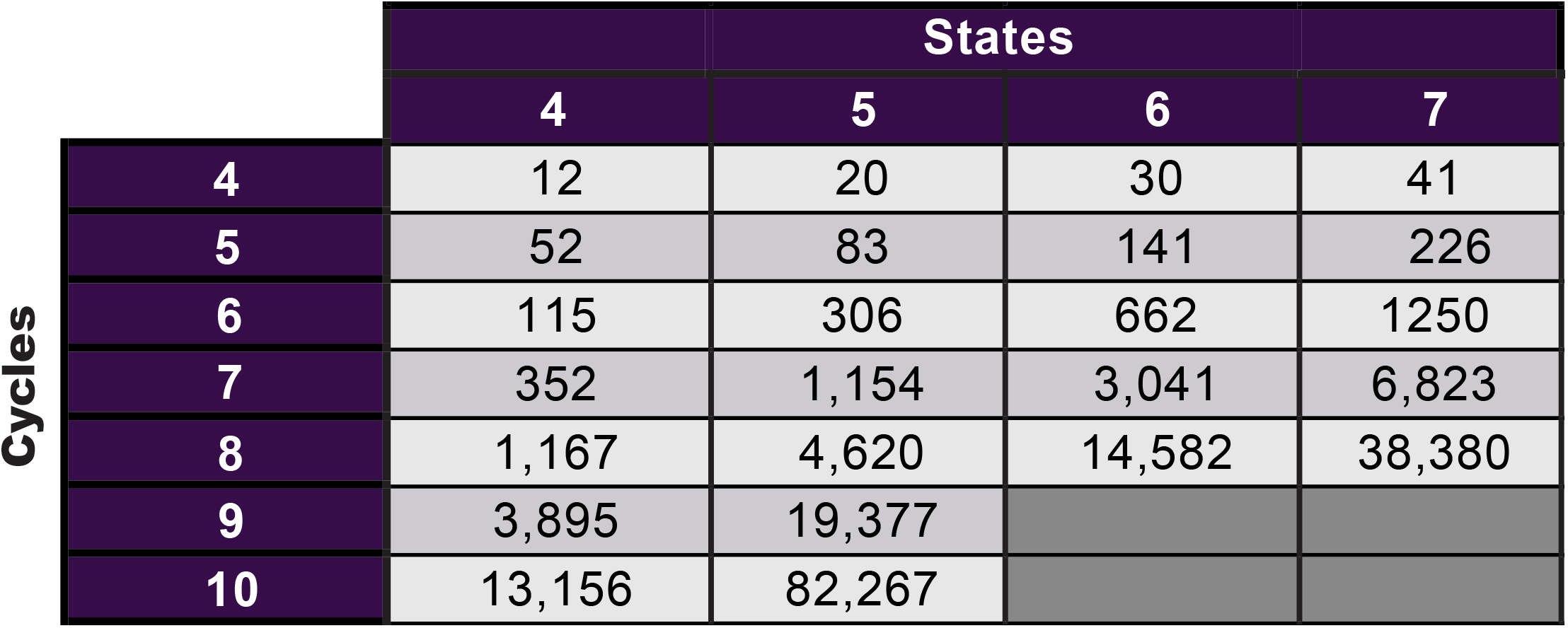
Hypercode plexities for several multi-state systems. Increasing either the number of readout cycles or the number of distinct identifiable optical states increases panel plexity exponentially.

|  |  |  | States |  |  |
| --- | --- | --- | --- | --- | --- |
|  |  | 4 | 5 | 6 | 7 |
| Cycles | 4 | 12 | 20 | 30 | 41 |
|  | 5 | 52 | 83 | 141 | 226 |
|  | 6 | 115 | 306 | 662 | 1250 |
|  | 7 | 352 | 1,154 | 3,041 | 6,823 |
|  | 8 | 1,167 | 4,620 | 14,582 | 38,380 |
|  | 9 | 3,895 | 19,377 |  |  |
|  | 10 | 13,156 | 82,267 |  |  |

**Extended Data Figure 1.**
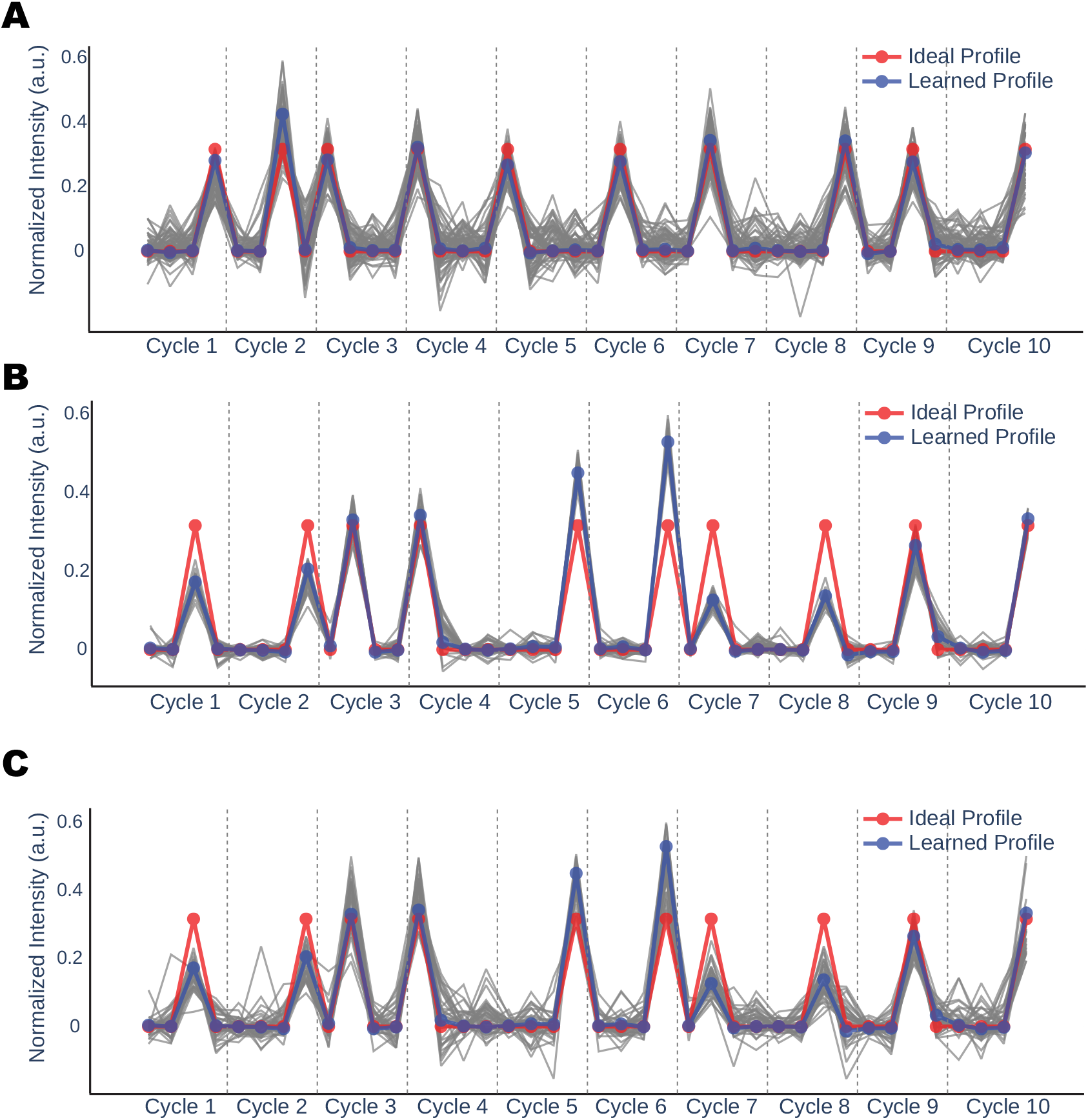
Example Hypercode intensity vectors and associated consensus Hypercode profiles. A. Representative intensity profiles for 100 RCPs (grey trace) in the 30th percentile (profile score <= 0.67) that decode to the Hypercode with median skew, 4311122434. The learned intensity profile (blue trace) and ideal intensity profile (red trace) are superimposed to reflect the systematic deviations in the measured intensity associated with this Hypercode. B. Representative intensity profiles for 100 RCPs (grey trace) in the 70th percentile (profile score >= 0.71) that decode to a Hypercode with high skew, 3421442334. C. Representative intensity profiles for 100 RCPs (grey trace) in the 30th percentile (profile score <= 0.65) that decode to a Hypercode with high skew, 3421442334.

**Extended Data Table 2.** Codespace sizes for a 4-state system. Increasing the codespace pairwise minimum Hamming distance cutoffs drastically reduces the pool plexity for all multi-flow systems.

|  |  | Pairwise Hamming distance minimum |  |  |
| --- | --- | --- | --- | --- |
|  |  | 3 | 4 | 5 |
| Cycles | 4 | 12 | 4 | 1 |
|  | 5 | 52 | 15 | 4 |
|  | 6 | 115 | 64 | 9 |
|  | 7 | 352 | 80 | 24 |
|  | 8 | 1,167 | 244 | 68 |
|  | 9 | 3,895 | 710 | 167 |
|  | 10 | 13,156 | 2,230 | 469 |

**Extended Data Table 3.** Decode performance with truncated intensity vectors. Decode error rates and the number of Hypercodes covered for both the 1K and 12K plex libraries. For nearly all the Hypercodes within the Plenoid library, excluding one cycle reduces the Hypercode pairwise Hamming distance from 3 to 2. Utilization of profile decoding compensates for profile skew and enables classification of most RCPs at <1/100,000 decode error rates.

|  | Cycles excluded<br>from analysis | Decode<br>Count | Decode<br>error rate | Hypercodes<br>covered |
| --- | --- | --- | --- | --- |
| 1K Plex | 0 | 1,959,438 | <1 in 100,000 | 1,032 counts $\geq 100$ |
| | 1 | 1,805,158 | <1 in 100,000 | 1,042 counts $\geq 100$ |
| 12K Plex | 0 | 1,036,565 | <1 in 100,000 | 11,293 counts $\geq 10$ |
| | 1 | 839,361 | <1 in 100,000 | 10,772 counts $\geq 10$ |

**Extended Data Table 4.** Small Variant Genotyping Performance. 209 variants were tested across 108 commercially acquired samples with indicated mean coverage and compared to truth data previously obtained using NGS. Numbers are mean ± standard deviation, when applicable.

| Panel Genotyping Metrics |  |
| --- | --- |
| Number of samples tested | 108 |
| Number of variants tested | 209 |
| Median coverage per locus | 1734 ± 335 |
| Call Rate across samples | 99.8 ± 0.3 % |
| Median concordance with 1000Genomes | 98.7 ± 0.9 % |

**Extended Data Figure 2.**
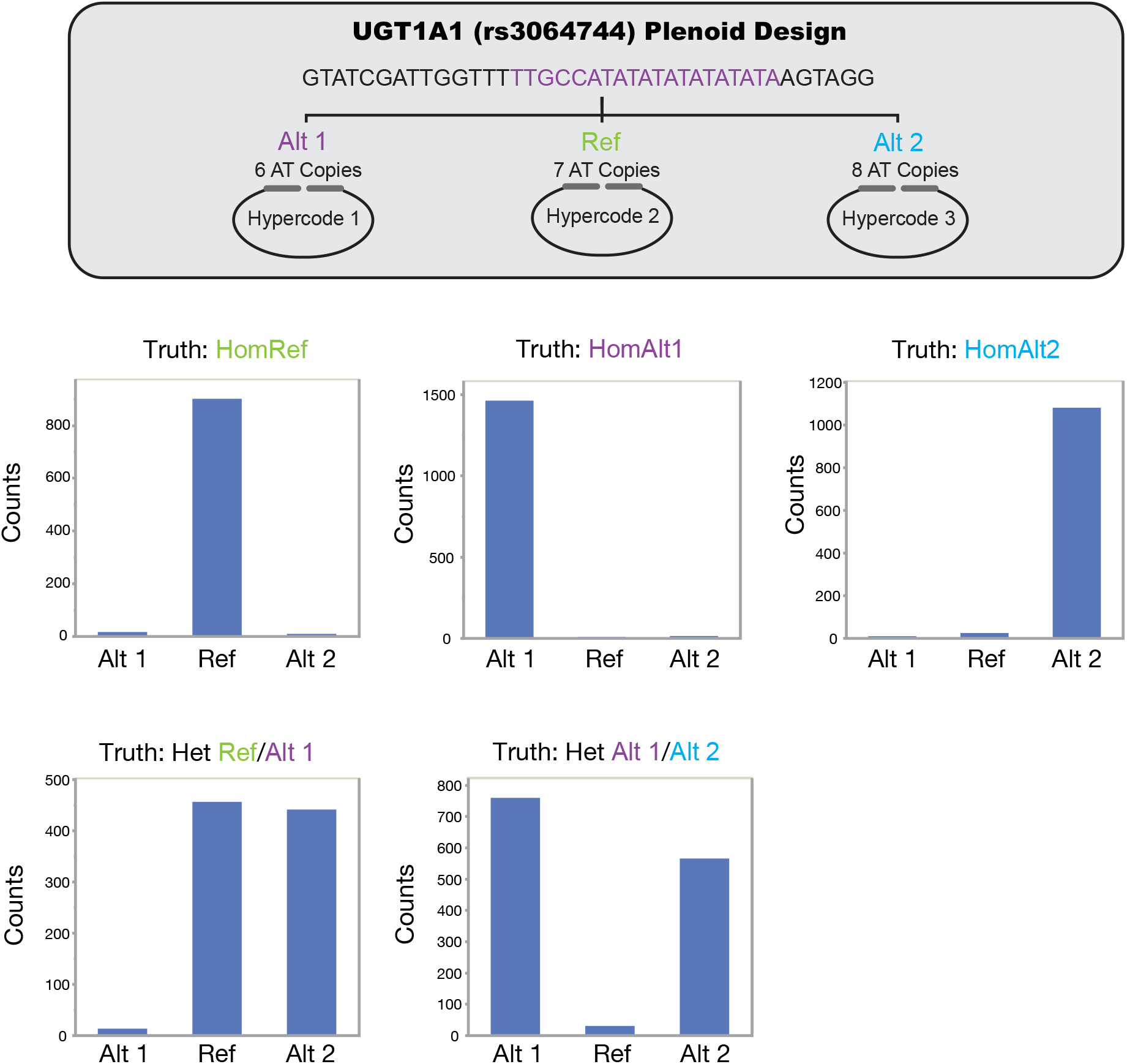
Germline genotyping of tandem repeats. The loss or gain of one ‘AT’ tandem repeat on the UGT1A1 gene can be accurately genotyped with good sensitivity for all alleles by using multiple Hypercodes and careful Plenoid design. Samples tested were HG00125 (HomRef), NA19771 (HomAlt 1), NA10831 (HomAlt 2), NA18499 (Het Ref/Alt 1), and NA19207 (Het Alt 1/Alt 2).

**Extended Data Table 5.** Accuracy of Copy Number Calling. Comparison of copy number calling using Hypercoding compared to the expected copy number truth as determined by NGS. Ratios are calculated for the number of correct calls made within each region for a particular copy number sample type.

| Region |  | CN = 1 |  | CN = 2 |  | CN = 3 |  | CN = 4 |
| --- | --- | --- | --- | --- | --- | --- | --- | --- |
|  | Intron1_Upstream | 100% (15/15) |  | 100% (29/29) |  | 100% (29/29) |  | 100% (6/6) |
|  | Intron1_Intron4 | 100% (19/19) |  | 100% (102/102) |  | 100% (29/29) |  | 100% (6/6) |
|  | Intron4_Exon9 | 100% (19/19) |  | 100% (102/102) |  | 100% (29/29) |  | 100% (6/6) |
|  | Exon9_Downstream | 100% (19/19) |  | 99.2% (118/119) |  | 100% (14/14) |  | 100% (4/4) |

**Extended Data Figure 3.**
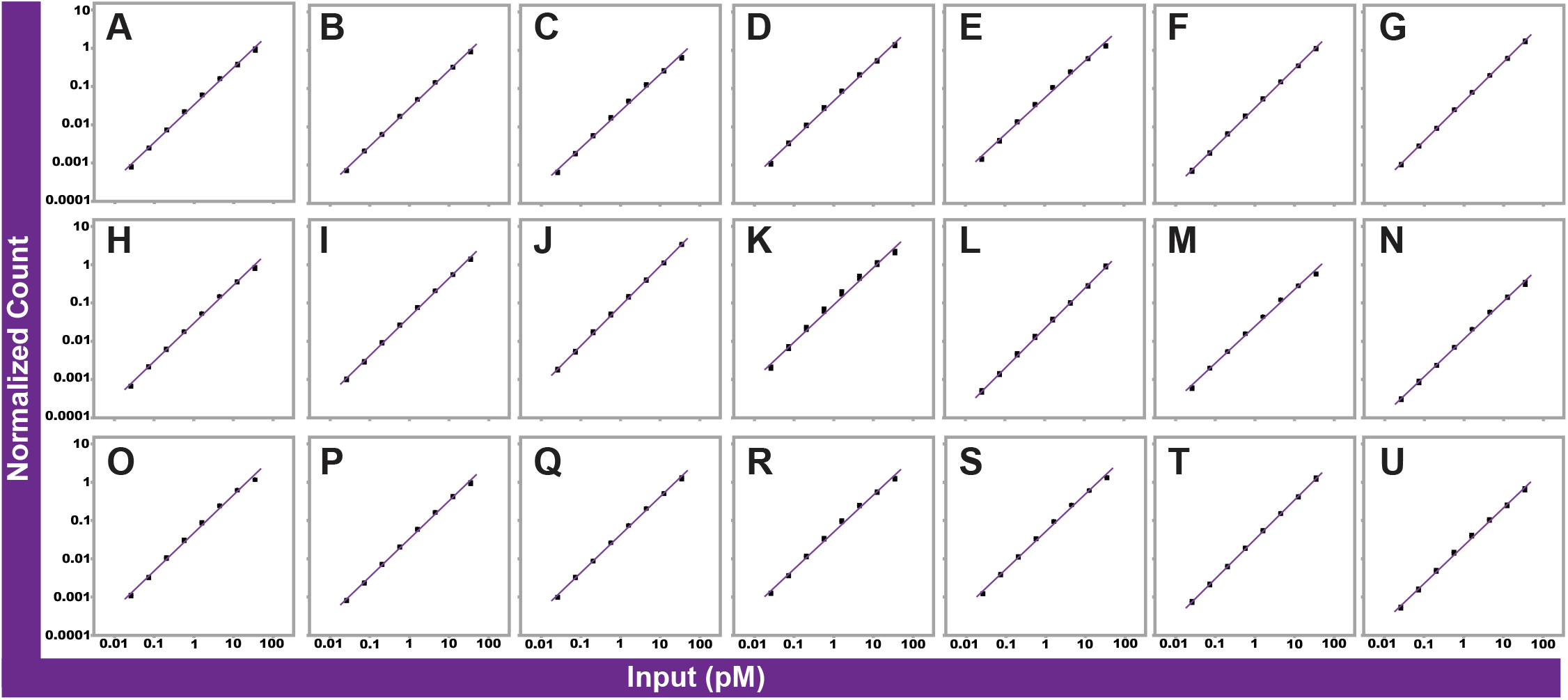
Linear response of individual targets over 3 log input. (A-U) Normalized counts as a function of input for 21 individual targets across 3 log of target input (0.03-30 pM) and associated linear fits. Normalization to internal calibrators compensates for well-to-well variability and the effects of surface saturation at high target input concentration.

**Extended Data Table 6.**
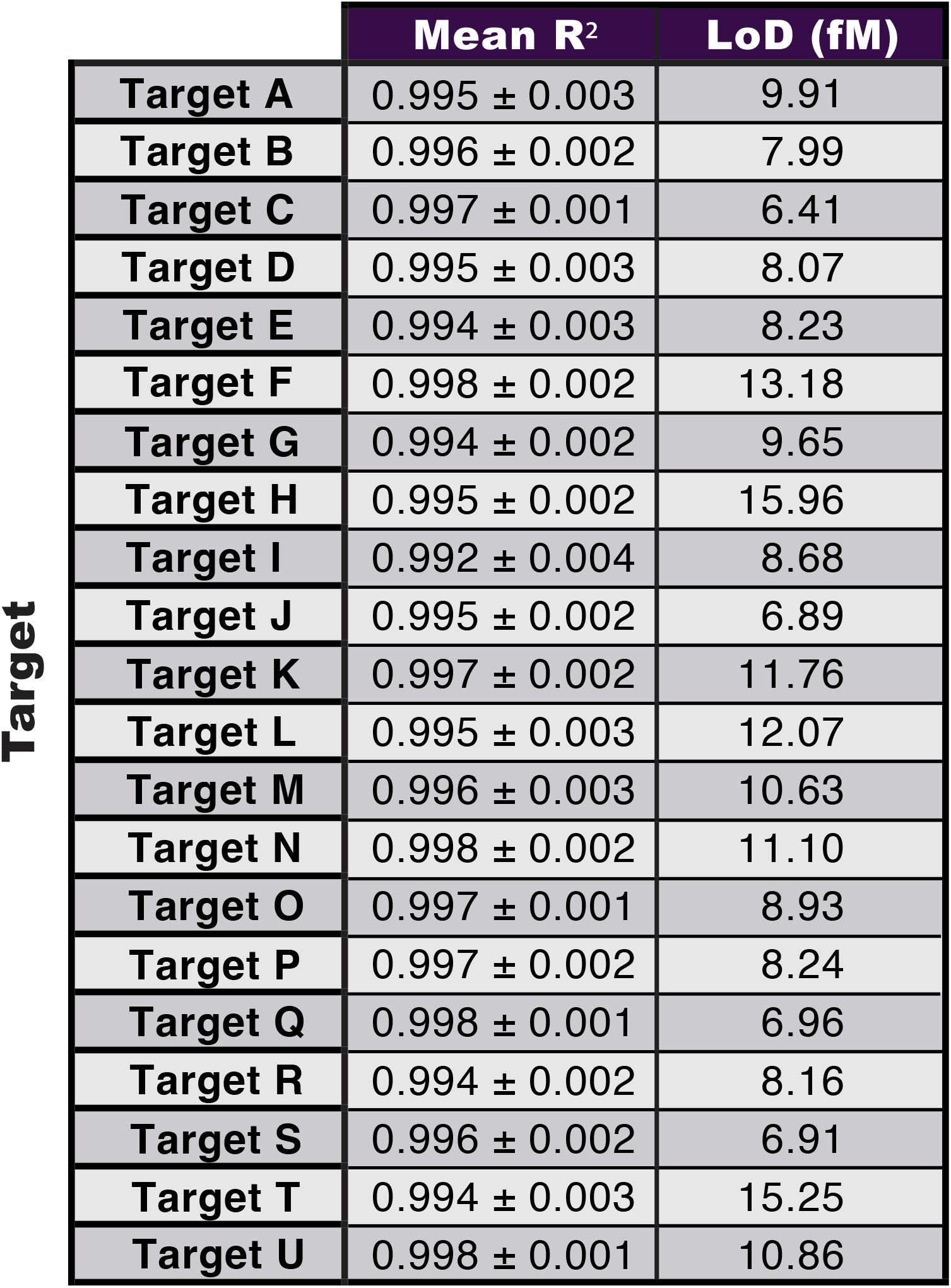
Accuracy and precision of quantification. Measured accuracy of fit for each target calibration in Extended Data Table 6. Calculated Limit of Detection (LoD) is also provided for each target using the previous fit equation along with the sample blank and lowest concentration.

| Target |  | Mean R<sup>2</sup> | LoD (fM) |
| --- | --- | --- | --- |
|  | Target A | 0.995 ± 0.003 | 9.91 |
|  | Target B | 0.996 ± 0.002 | 7.99 |
|  | Target C | 0.997 ± 0.001 | 6.41 |
|  | Target D | 0.995 ± 0.003 | 8.07 |
|  | Target E | 0.994 ± 0.003 | 8.23 |
|  | Target F | 0.998 ± 0.002 | 13.18 |
|  | Target G | 0.994 ± 0.002 | 9.65 |
|  | Target H | 0.995 ± 0.002 | 15.96 |
|  | Target I | 0.992 ± 0.004 | 8.68 |
|  | Target J | 0.995 ± 0.002 | 6.89 |
|  | Target K | 0.997 ± 0.002 | 11.76 |
|  | Target L | 0.995 ± 0.003 | 12.07 |
|  | Target M | 0.996 ± 0.003 | 10.63 |
|  | Target N | 0.998 ± 0.002 | 11.10 |
|  | Target O | 0.997 ± 0.001 | 8.93 |
|  | Target P | 0.997 ± 0.002 | 8.24 |
|  | Target Q | 0.998 ± 0.001 | 6.96 |
|  | Target R | 0.994 ± 0.002 | 8.16 |
|  | Target S | 0.996 ± 0.002 | 6.91 |
|  | Target T | 0.994 ± 0.003 | 15.25 |
|  | Target U | 0.998 ± 0.001 | 10.86 |

**Extended Data Table 7.** Accuracy and precision of quantification. Percent error and inter-assay (plate-to-plate) percent CV (mean ± standard deviation) for the calculated concentration of each target in the 7 pools for high (30 pM), medium (1 pM), and low (0.03 pM) input concentrations. The average percent error across the entire concentration range was 18.8± 0.9%.

|  | 0.03 pM |  | 1 pM |  | 30 pM |  |
| --- | --- | --- | --- | --- | --- | --- |
|  | % Error | % CV | % Error | % CV | % Error | % CV |
| Target A | 25.5 | 25.4 | 14.5 | 15.9 | 9.0 | 11.9 |
| Target B | 21.8 | 25.3 | 16.6 | 17.8 | 13.0 | 9.1 |
| Target C | 23.7 | 29.2 | 22.8 | 8.1 | 11.2 | 12.2 |
| Target D | 20.4 | 23.6 | 12.5 | 16.0 | 10.5 | 9.8 |
| Target E | 16.7 | 21.6 | 15.8 | 14.8 | 12.5 | 11.6 |
| Target F | 21.5 | 23.5 | 21.2 | 12.7 | 15.3 | 9.4 |
| Target G | 41.8 | 25.7 | 17.7 | 14.5 | 12.6 | 15.1 |
| Target H | 64.1 | 26.9 | 11.6 | 9.9 | 15.6 | 13.0 |
| Target I | 29.8 | 29.5 | 23.5 | 13.6 | 12.4 | 15.6 |
| Target J | 68.8 | 29.2 | 28.9 | 10.9 | 10.2 | 10.9 |
| Target K | 12.7 | 18.2 | 10.2 | 11.7 | 7.4 | 7.3 |
| Target L | 60.5 | 22.6 | 10.0 | 12.4 | 24.2 | 12.3 |
| Target M | 25.5 | 34.3 | 8.2 | 11.4 | 25.9 | 15.5 |
| Target N | 23.0 | 32.0 | 13.0 | 19.8 | 11.7 | 17.4 |
| Target O | 13.0 | 19.9 | 29.1 | 15.4 | 7.0 | 7.0 |
| Target P | 19.8 | 25.5 | 11.4 | 7.6 | 11.0 | 10.0 |
| Target Q | 177.3 | 26.3 | 10.1 | 10.0 | 12.0 | 11.3 |
| Target R | 17.7 | 16.1 | 12.2 | 13.4 | 16.1 | 16.4 |
| Target S | 34.0 | 33.1 | 13.2 | 17.3 | 12.1 | 10.8 |
| Target T | 19.9 | 20.2 | 11.2 | 14.0 | 14.1 | 12.6 |
| Target U | 154.0 | 22.7 | 13.1 | 13.7 | 12.4 | 10.6 |
|  | 42.5±44.2 | 25.3±4.8 | 15.6±6.12 | 13.4±3.1 | 13.1±4.6 | 11.9±2.8 |

